# Improving the solubility of pseudo-hydrophobic Alzheimer’s Disease medicinal chemicals through co-crystal formulation

**DOI:** 10.1101/2023.04.25.538327

**Authors:** Tse A, Janilkarn-Urena I, J Lin, X Chang, C Efthymiou, A Idrissova, M Zhang, Williams CK, S Magaki, Vinters HV, Davies DL, T Gonen, Gukasyan HJ, Seidler PM

## Abstract

Natural products are ligands and potential inhibitors of Alzheimer’s disease (AD) tau. Dihydromyricetin (DHM) is a CNS active natural product. Despite having signature polyphenolic character, DHM is ostensibly hydrophobic owing to intermolecular hydrogen bonds that shield hydrophilic phenols. Our research shows DHM becomes ionized at near-neutral pH allowing formulation of salts with transformed solubility. The MicroED co-crystal structure with trolamine reveals DHM salts as metastable solids with unlocked hydrogen bonding and a thermodynamic bent to solubilize in water. All salt formulations show better inhibitory activity against AD tau than the non-salt form, with efficacies correlating to enhanced solubilities. These results underscore the role of structural chemistry in guiding selection of solubilizing agents for chemical formulation. We propose DHM salts are appropriate formulations for research as dietary supplements to promote healthy aging by combating protein misfolding. Additionally, DHM is a suitable lead for medicinal chemistry and possible development of CNS pharmaceuticals.

## Introduction

Solubility is a crucial factor affecting the efficacy of pharmaceuticals and other bioactive compounds such as vitamins, natural products, and dietary supplements. The degree of absorption is directly proportional to the solubility of a compound, although the physical and chemical mechanisms underlying solubility are largely theoretical. Compounds that deviate from ideality pose a challenge to scientists seeking to troubleshoot and develop rational strategies to overcome poor chemical solubility. Approximately 40% of new pharmaceutical chemicals developed by the industry are described as practically insoluble in water^1^ and nearly 90% of new drugs are classified accordingly as Class II/IV in the Biopharmaceutics Classification System (BCS)^2^ partly due to low solubility. Salt formulations are a common strategy for preparing stable, safe, and bioavailable dosage forms in the pharmaceutical industry.^3^ However, predicting optimal salt forms and how they will function remains challenging.

In the present work, we investigated the solubility, activity, and Microcrystal electron diffraction (MicroED)^4, 5^ structure of salt formulations of the natural product dihydromyricetin (DHM), a brain-permeable flavonoid with high chemical similarity to a known in vitro tau inhibitor, EGCG^6^. DHM is a plant-derived polyphenol with antioxidant^7, 8^ and anti-inflammatory^9–11^ activity and potential benefits in ameliorating dyslipidemia^7, 9, 12, 13^ and alcohol intoxication^14, 15^. The CNS effects of DHM are attributed to increased GABAergic transmission and synaptic functioning, reduced neuroinflammation, and restoration of redox imbalances in neurons through improved mitochondrial function.^16–18^ Despite its rich phenolic character, DHM behaves as a hydrophobic substance with an aqueous solubility of only ∼0.4 mg/ml. While hydrophobic character of DHM allows for permeability to the CNS, poor water solubility impedes dosing and intestinal absorption.^19–22^ Water-soluble formulations of DHM could overcome dosing issues by enabling delivery of higher dosage concentrations to enable increased absorption.

Like salts, co-crystal solid forms of organic molecules typically display a defined stoichiometry of solid components (as opposed to liquid residues) found within their crystalline lattice. While they allow for higher dissolved concentrations in bulk solution than their respective free acid or base (non-ionized) form, co-crystal lattice constituents distinguish themselves from salts with nonionic interactions. Suitable counterions are found on the FDA-approved list of Inactive Ingredients Database.^23^ While natural flavonoids and DHM are generally regarded as safe (GRAS), feasibility of isolating them as salts or co-crystals with suitable counterions that improve poor physicochemical properties (e.g., low solubility, instability or hygroscopicity) has not been widely explored.^24^ Several examples of other pharmaceutical co-crystal formulations have been commercialized, e.g., Entresto® ^25, 26^ and Steglatro® ^27^, with the latter being a medicinal chemistry derived analog of phlorizin (i.e. another flavonoid) ^28^. In this study, we aimed to improve DHM solubility by salt/co-crystal formulation and explored DHM’s in vitro activity to disrupt prionogenic seeding by AD tau tangles.

With millions aging into elevated risk for AD, an elderly population that is expected to double by 2050, and knowledge that AD pathophysiology accrues over the course of decades, there is pressing need for comprehensive public health action. The U.S. Department of Health and Human Services outlined a “National Plan to Address Alzheimer’s Disease” aiming to “Prevent” and “Treat” AD by 2025 by dual focus on supporting healthier aging and developing pharmaceutical therapies to manage and cure AD.^29^ Epidemiological research suggests lifestyle and environmental factors convolve with genetics and aging to affect timing of molecular transformations that lead to AD. This, and realization that AD manifests after decades in the making inspires research to delay AD onset by managing lifestyle interventions in the decades that precede AD dementia. Current prevention strategies being investigated includes dietary and exercise lifestyle adaptions, and attention to cardiovascular health and physical wellbeing. Lifestyle adaptions that support healthier aging paired with biologically active chemicals, such as pharmaceutics and dietary supplements could provide a tiered and currently missing framework to manage aging and AD by leveraging early chemical intervention.

Tau tangles are thought to kill neurons in AD and promote dozens of other neurodegenerative tauopathies, so suppressing tau spreading is seen as a top interest therapeutic mechanism of action. Despite having outstanding in vitro inhibitory activity towards seeding by AD tau, EGCG is limited by poor brain permeation. Based on the CryoEM structure of AD tau fibrils bound with EGCG and congeneric substructure realized in DHM, we identified DHM as a possible alternative natural product and brain-permeable tau inhibitor, which spurred our investigations of its bioactivity towards AD tau.

## Results

The polyphenol DHM has similar chemical structure to EGCG, a known polyphenolic ligand and inhibitor of tau (Fig. 1A). Despite rich phenolic character, DHM is notoriously hydrophobic with a reported water solubility of ∼0.2-0.4 mg/ml and enhanced ability to partition to the CNS compared to other polyphenols. Therefore, we evaluated the in silico predicted binding of DHM to AD tau paired helical filaments (PHFs) using CB-Dock^30^, a suite that incorporates cavity detection with AutoDock Vina. Using an AD tau PHF co-cryoEM structure with EGCG as a starting model, we evaluated the most likely binding cavities for DHM. The top-scoring predicted DHM binding cavity matched the primary EGCG binding volume formerly determined by CryoEM (Fig. 1A), although the DHM binding pose output by CB-Dock differs from models we generated manually by superimposing DHM with EGCG from the liganded CryoEM structure (Supplemental Fig. 1). Both modeling approaches predict interactions with Lys340, although models generated by CB-Dock orient the π orbitals of aromatic moieties of DHM perpendicular to the fibril axis whereas manually generated models orient stacks of aromatic moieties parallel to strand forming β sheets of the fibril. Additional possible DHM binding sites detected by CB-Dock include cavities labeled Sites 2 and 3 (Fig. 1A). However, both Sites 2 and 3 scored lower by AutoDock Vina. Site 2 is lined by histidine and lysine residues, which could support hydrogen bonding interactions with DHM, similar to the EGCG binding site of Tau PHFs. Site 3 is created by a sharp turn at a histidine and glycine interface and scores significantly lower favorably compared to Sites 1 and 2.

**Figure 1.**
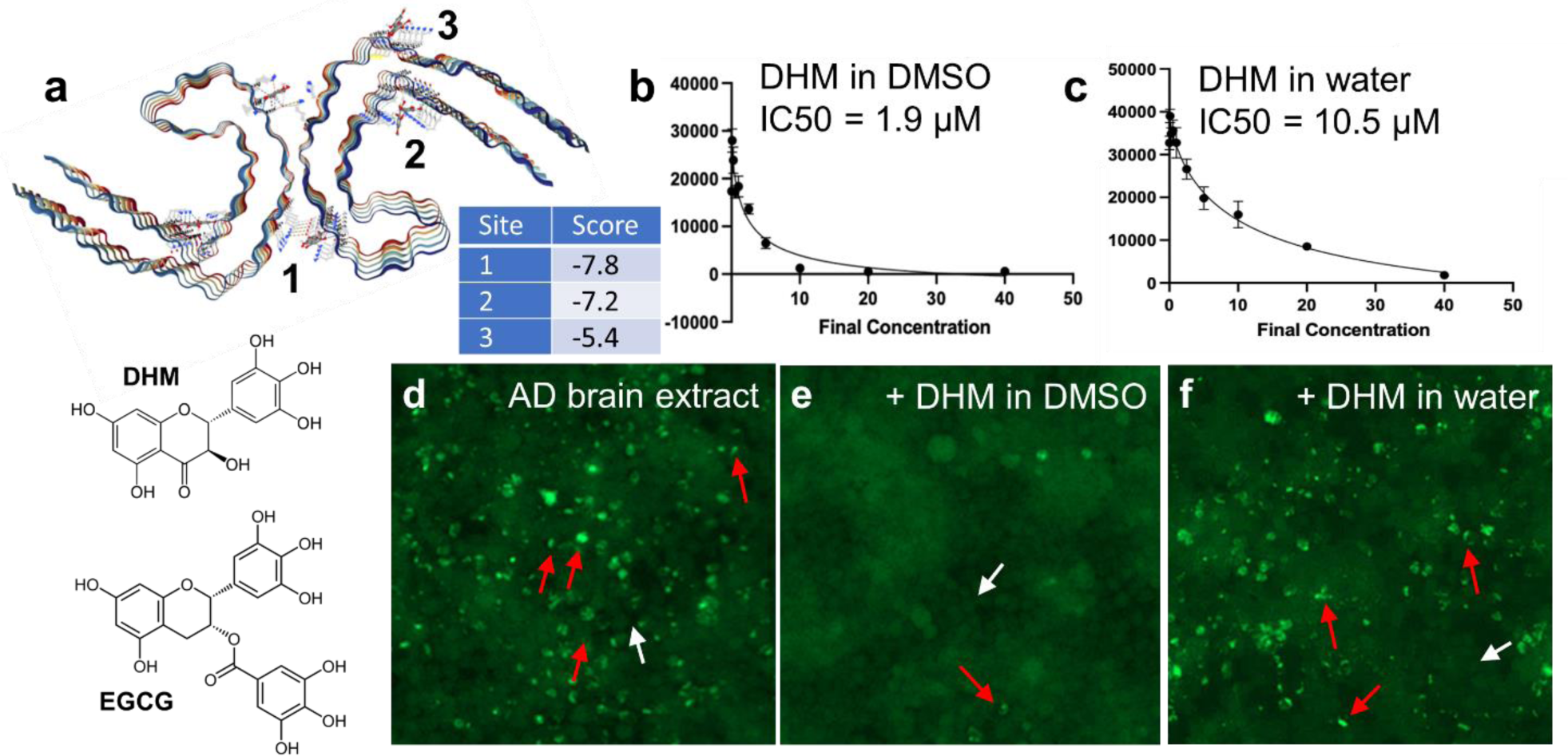
(a) DHM binding sites on AD tau fibrils predicted by CB-Dock. AutoDock Vina docking scores are shown in the embedded table. Chemical structures comparing DHM and EGCG are shown below the CB-Dock model (b-c) Dose dependent seeding inhibition measured by transfecting AD crude brain homogenates in HEK293 tau biosensor cells that stably express

P301S 4R1N tau fused to YFP. Seeding inhibition was determined by counting the number of fluorescent puncta as a function of inhibitor concentration. IC50 values were calculated by nonlinear curve fitting from dose–response plots. Experiments in b were performed by pre-incubating AD crude brain homogenates with DHM dissolved in DMSO. Experiments in c were performed identically except using the soluble fraction of DHM dissolved in water following centrifugation at 8,000 rpm. (d-f) Representative images from tau biosensor cells transfected with AD crude brain homogenate (d), or identically treated cells following pre-incubation of AD crude brain homogenate with DHM (10 mM final concentration on cells). Representative tau-4R1N cells containing aggregated tau puncta are marked with red arrows, and cells without are marked with white arrows.

We tested the in vitro activity of DHM to inhibit prionogenic seeding by AD tau (Fig. 1b-f) since modeling suggests DHM can bind to AD tau fibrils in a manner similar to EGCG. Transfecting AD crude brain homogenates in tau biosensor cells, which express an aggregation-prone fragment of tau fused with YFP reporter, produces a punctated cellular phenotype, shown in Fig. 1d that results from the aggregation of intracellular tau that is seeded by tau from AD brain homogenate. Pre-treating AD crude brain homogenates with DHM dissolved in DMSO inhibited seeding in a dose-dependent manner with an IC50 of 1.9 µM (Fig. 1b). In comparison, DHM that was dissolved in water elicited less potent inhibition with an apparent IC50 of 10.5 µM (Fig. 1c). These results were consistent with limiting solubilities observed for DHM stock solutions prepared in water (Supplementary Fig. 2). We observed that 10 mM master stock solutions of DHM prepared in water were unstable, resulting in DHM precipitation and eventual equilibration to a stock solution with actual measured concentrations of 0.65 mg/ml (2 mM). In contrast, stock solutions prepared in DMSO remained soluble. Our observation that only about one-fifth of DHM remained dissolved in aqueous solutions after clearing insoluble matter by centrifugation may account for the lowered measured IC50 of DHM solutions prepared in water. These data illustrate two important points: (1) DHM effectively inhibits seeding by AD tau, particularly when solubilized in nonpolar solvents, and (2) the inhibitory power of DHM is limited by poor aqueous solubility.

Compounds with limiting aqueous solubilities exhibit low bioavailability and compromised bioactivity. Therefore, we investigated the physicochemical factors underlying the poor aqueous solubility of DHM with the goal of developing formulations to overcome the solubility limitation. An analysis of pKa shown in Fig. 2a reveals plausible ionization states of DHM. A first acid dissociation event is predicted by ADMET Predictor (Simulations Plus, Inc., Lancaster, CA) to occur between pH 7.05 ↔ 7.83, with DHM transitioning from neutral to a negatively charged ion. Four primary sites of possible ionization shown in Fig. 2a are predicted in varying ratios.

**Figure 2.**
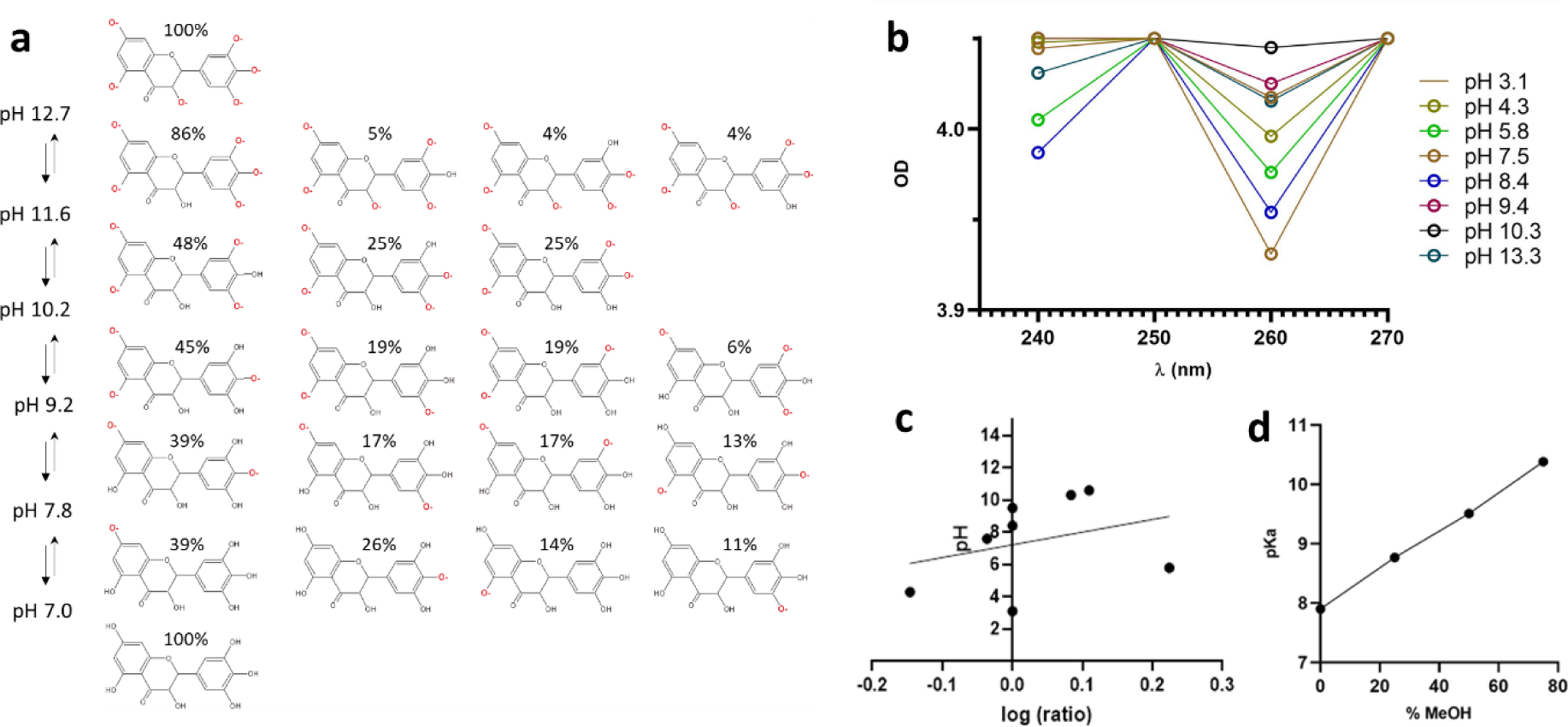
Predicted and experimental pKa’s of DHM. (a) Ionizations occurring from neutral to anionic states produced using ADMET Predictor™. Predicted proportions of respective ionized states are indicated. Microstates for multiply ionized phenols are shown as a function of increasing pH. (b and c) UV-VIS spectroscopic pKa determination of DHM determined at pH values ranging from pH 3.1-13.3. (d) NaOH potentiometric titration curves with MeOH revealing an equivalence point at ∼pH 7.9.

The predicted near neutral pKa for DHM was verified experimentally by two methods: potentiometric titration and UV-Vis spectroscopy using a modified method published^31^. Experimental pKa values presented in Figure 2 b-c were obtained by plotting the absorbance ratio of DHM at 240 and 260 nm across a range of pH values (3.1-13.3) in various buffer solutions (as listed in Supplementary Table 1). The determined pKa, pH 7.9, was taken as the Y intercept of the line of best fit of the log ratio (Fig. 2c). To confirm the accuracy of the pKa value obtained via this method, DHM was subjected to potentiometric titration with sodium hydroxide in methanol, and the resulting pKa value was found to be in agreement with the pH 7.9 value (as depicted in Figure 2d).

We investigated the solubility-enhancing properties of the following five complexes with DHM: triethanolamine (TEA), sodium hydroxide (NaOH), TRIS Base (2-Amino-2-(hydroxymethyl)-1,3-propanediol), L-lysine, and calcium hydroxide (CaOH_2_), each combined with DHM in one of four solvents (ethanol, methanol, isopropanol, or acetone) and rendered by slow evaporation. New salt or co-crystal forms of DHM were made by setting up a reaction in a solvent system which allows DHM and counterion to fully dissolve and facilitate proton transfer from DHM (proton donor) to a base (proton acceptor). We set up a counterion screen and produced several slurries, using various pharmaceutically relevant counterions in different solvent systems (Supplementary Table 2 and Supplementary Figure 3). Five of the combinations yielded powders, which were taken as potential salt formulations: DHM-TEA prepared from ethanol, DHM-Tris prepared from isopropanol, DHM-Lysine prepared from water, and two DHM-Ca formulations: DHM-Ca(b) and DHM-Ca(c), prepared form methanol and 2-propanol, respectively.

The five salt forms were tested for tau inhibitor activity by dissolving in water to a target concentration of 10 mM. As shown in the schematic in Fig. 3a outlining our workflow, insoluble DHM was removed from concentrated stocks by centrifugation to yield supernatants that were tested for tau inhibitor activity. We reasoned that aqueously soluble fractions of DHM salts would exhibit tau inhibitor activities with potencies that approached DHM dissolved in DMSO. DHM-TEA, -Lysine, -Tris, -Ca all exhibited inhibitor activity compared with DHM (Fig 3b-e). DHM-TEA, -DHM-Ca were of greatest improved tau inhibitor potencies. The IC50s of DHM-TEA and DHM-Ca(b) were 2.6 and 0.87 µM, respectively, (Fig. 3d and e), similar to the IC50 we measured for DHM dissolved in DMSO.

**Figure 3.**
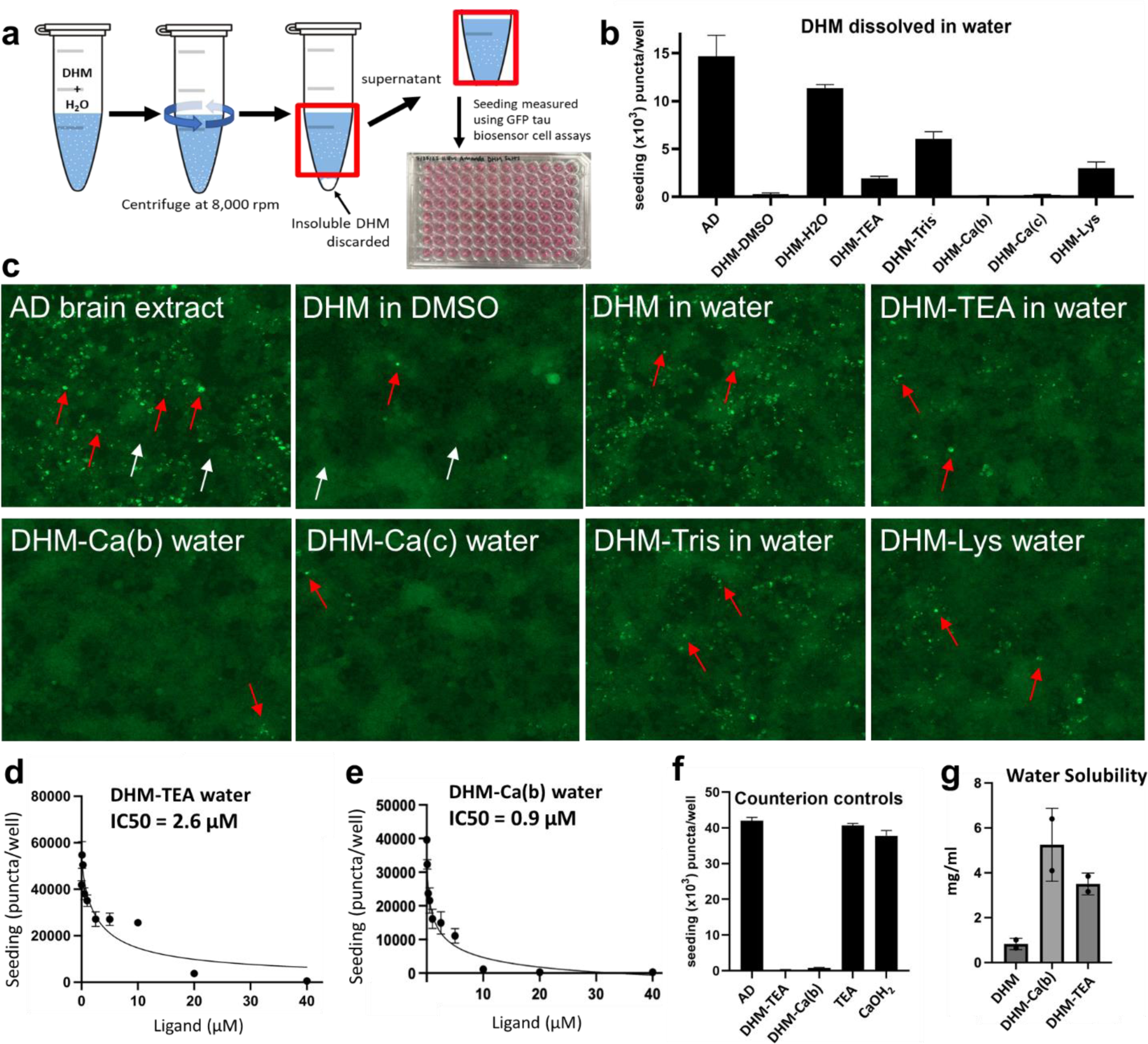
(a) Schematic showing experimental design of experiments in Figure 3. Co-crystalline DHM formulations were dissolved in water to a target 10 mM concentration. Insoluble matter was removed by centrifugation as indicated. Supernatants containing water solubilized DHM were assayed for tau inhibitor activity. (b) Seeding inhibition measured by quantifying the number of fluorescent puncta as a function of indicated inhibitor. AD sample is AD brain homogenate without added inhibitor. Error bars represent standard deviations of triplicate measures. (c) Representative images from b of tau biosensor cells seeded by transfection by AD crude brain homogenates following pre-treatment with DHM or salt co-crystals. Inhibitors were added to a final concentration of 10 µM on cells. Example puncta are shown by red arrows, examples of non-seeded cells shown with white arrows. (d and e) IC50 dose dependent inhibition of puncta following pre-treatment with DHM-TEA and -Ca(b). (f) Seeding inhibition measured for DHM-TEA and -Ca(b), or TEA and -CaOH_2_. (g) Solubility determination of DHM, DHM-TEA and DHM-Ca(b) in water, expressed in mg/ml.

As a control to ensure counterions themselves do not contribute to the measured tau inhibitory activities, we measured tau seeding after pre-incubating AD brain homogenates with DHM salts and counterions. As shown in Fig 3f, neither TEA nor CaOH_2_ counterion had inhibitory activity towards tau seeding by crude AD brain homogenates, confirming that the DHM inhibitor activity is attributable to improved solubility of DHM and not the result of synergistic inhibition by accompanying counterions. Also supporting this are data in Supplementary Figure 4 showing DHM salts DHM-TEA and -DHM-Ca(b) possess IC50s of ∼1 µM similar to DHM, indicating that the counterions themselves do not enhance tau inhibitor activity beyond what is possible for DHM itself dissolved at the target concentration. Thus, counterions improve inhibitory activity of DHM by effecting solubility. Experimental solubilities of DHM-TEA and -DHM-Ca shown in Fig 3g reveals 4-6 fold improved water solubility, consistent with the fold improvements in IC50 that were seen for these salts.

Since DHM shares a similar structure to EGCG, we tested the hypothesis that DHM inhibits seeding by a similar mechanism by AD tau disaggregating fibrils. AD tau fibril disaggregation by DHM was measured by quantitative electron microscopy (qEM). As shown in micrographs in Fig. 4, incubation with EGCG reduces the average number of AD tau fibrils observed by negative-stain EM imaging by 70-80%. DHM-TEA and -Ca(b) both reduced fibril density by 50%, from 300 to 160 fibrils after 48 hrs incubation with DHM-TEA and from 227 total fibrils to 111 fibrils after 48 hrs incubation with DHM-Ca(b). These data suggest fibril disaggregation is one plausible mechanism of inhibition, although they leave open the possibility that DHM binding to AD tau is inhibitory towards tau seeding since fibril disaggregation occurs to a lesser extent compared to EGCG. Further supporting the possibility that ligand binding exerts inhibitory effects absent of fibril disaggregation are data showing that inhibitor pre-incubation is not needed to realize seeding inhibition (Supplementary Figure 5). These data showing that seeding inhibition by DHM occurs despite some fraction of remaining fibrils in solution thus suggest it is possible that ligand binding itself suppresses seeding, at least to a partial degree.

**Figure 4.**
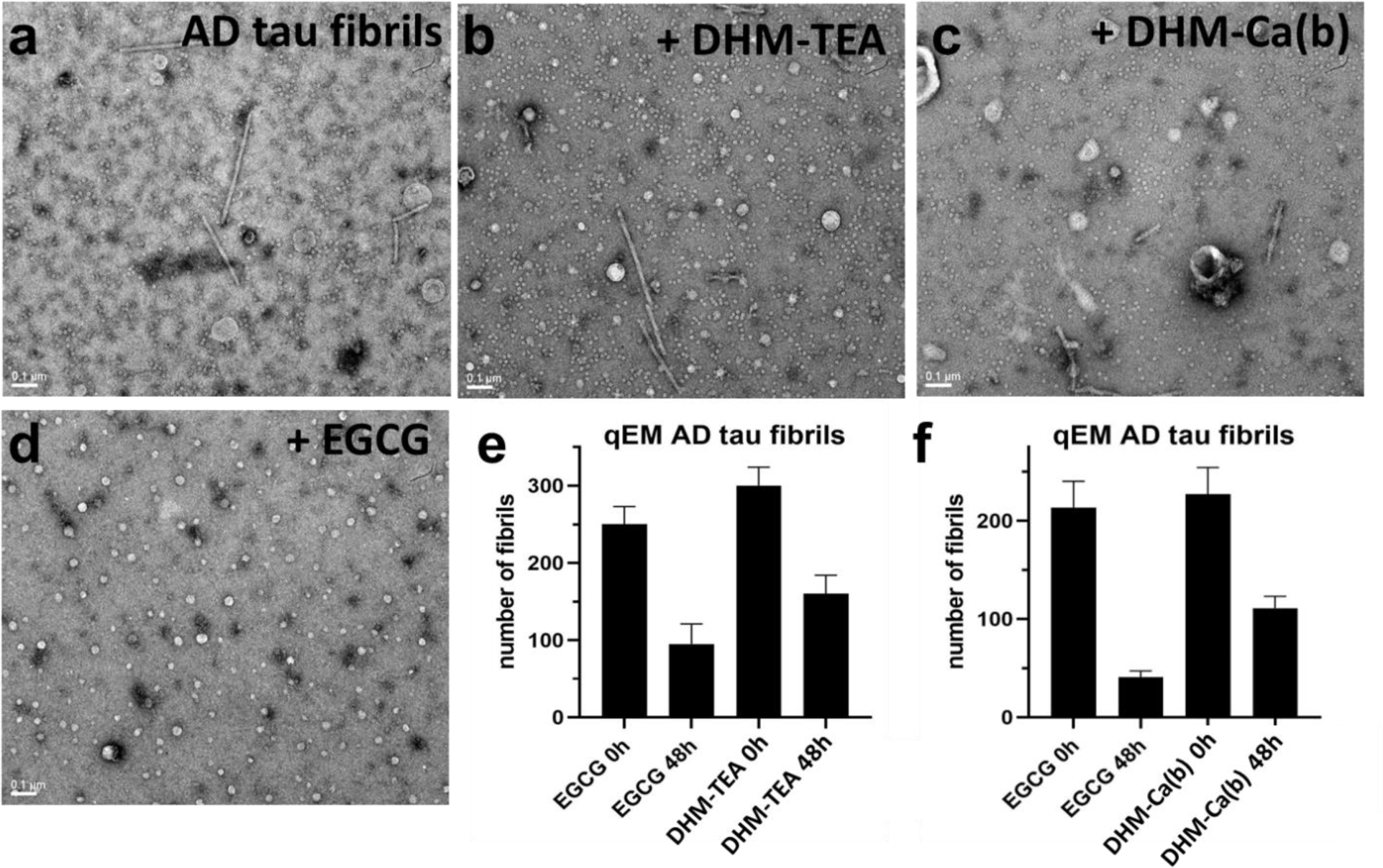
(A) DHM-mediated AD tau fibril disaggregation, measured by qEM. Fibrils were counted from N=90 randomly acquired EM images, which were split three ways and quantified in triplicate. Columns show numbers of fibrils counted as a function of inhibitor pre-incubation time, as indicated. Error bars represent standard deviations. (B-D) Example EM images used from qEM after 48 hr inhibitor incubation.

The mechanism of enhanced solubility of DHM salt was investigated by structural chemistry. DHM-TEA and -Ca exhibited X-ray powder diffraction indicating crystallinity (Fig. 5a and Supplementary Figures 6 and 7), although only DHM-TEA produced suitable diffraction by MicroED to yield an atomic structure. TEA is seen in the 0.74 Å resolution MicroED structure in a 1:1 molar ratio positioned 2.6 Å from the O2 of the phenol of the resorcinol that is predicted to first become ionized (Fig. 5b and Fig. 2a). Although crystals of DHM-Ca(b) also diffracted, the diffraction power was insufficient to enable atomic structure determination. It is possible the tridentate nature of triethanolamine reinforced the crystal lattice enabled stronger diffraction and structure elucidation compared with DHM-Ca salts, which were crystalline but exhibited weaker diffraction.

**Figure 5.**
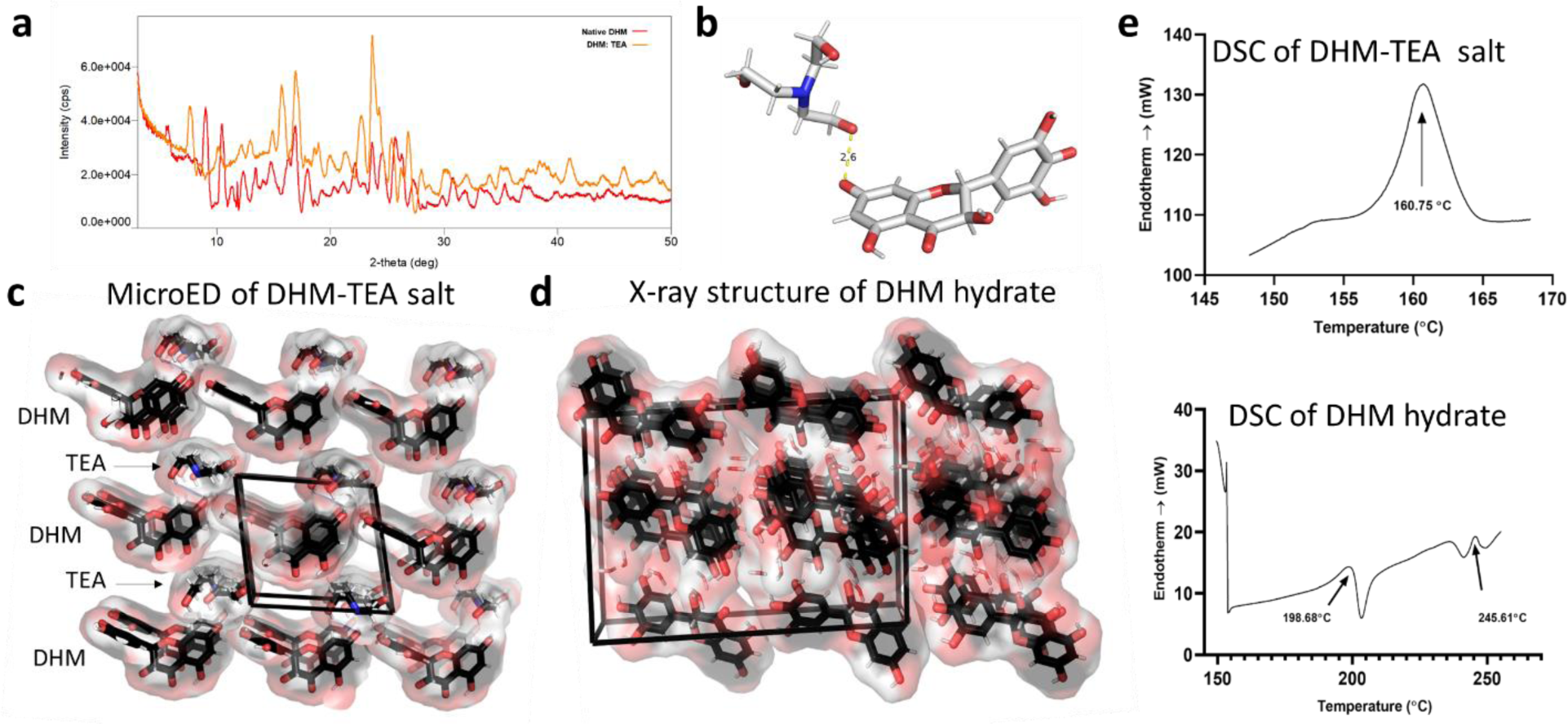
(a) X-Ray powder diffractogram comparing native dihydromyricetin (DHM) shown in red, overlayed against co-former dihydromyricetin with triethanolamine shown in orange. (b) Asymmetric unit of DHM complex with TEA determined by MicroED. (c) Crystal lattice of DHM-TEA salt. Note solvent channels created by the intercalation of TEA in the lattice between DHM molecules. (d) X-ray structure of DHM dihydrate^32^. Differential scanning calorimetry thermographs of DHM-TEA and DHM dihydrate.

Comparing the DHM-TEA dihydrate^32^ lattices in (Fig. 5c and d) reveals large solvent channels and high solvent content in the TEA lattice. Analysis of porosity using CrystalMaker reveals the DHM dihydrate unit cell is 20.6% filled space with 79.4% void. Co-crystallizing with TEA alters DHM packing leading to a fractured network of intermolecular phenol-phenol H-bonds that otherwise distinguishes the dihydrate lattice. As a result, the DHM-TEA lattice is relatively porous with void volume increased to 83.3% of the unit cell volume. The unit cell volume of DHM dihydrate, 2899.1 Å^3^, has a low solvent content described by a Matthew Coefficient (V_M_) of 1.02 Å^3^/Da, which is typical of small molecule crystals. The V_M_ of DHM with a unit cell volume of 971.5 Å^3^ is 2.07 Å^3^/Da assuming a mass of 469 da, which is atypically large for a small molecule crystals and more closely mirrors the V_M_ of protein crystals which are entrenched with large solvent channels enabling high solvent content^33^. Large solvent channels permeating the lattice of DHM-TEA lattice are seen in Fig. 5c and 6a, and likely enable water molecules to infuse and hydrate molecules of DHM more readily than for the DHM dihydrate. Correspondingly, DHM-TEA exhibit a cooperative transition at lower temperature suggesting DHM-TEA is less thermodynamically stable than DHM dihydrate (Fig. 5e).

**Figure 6.**
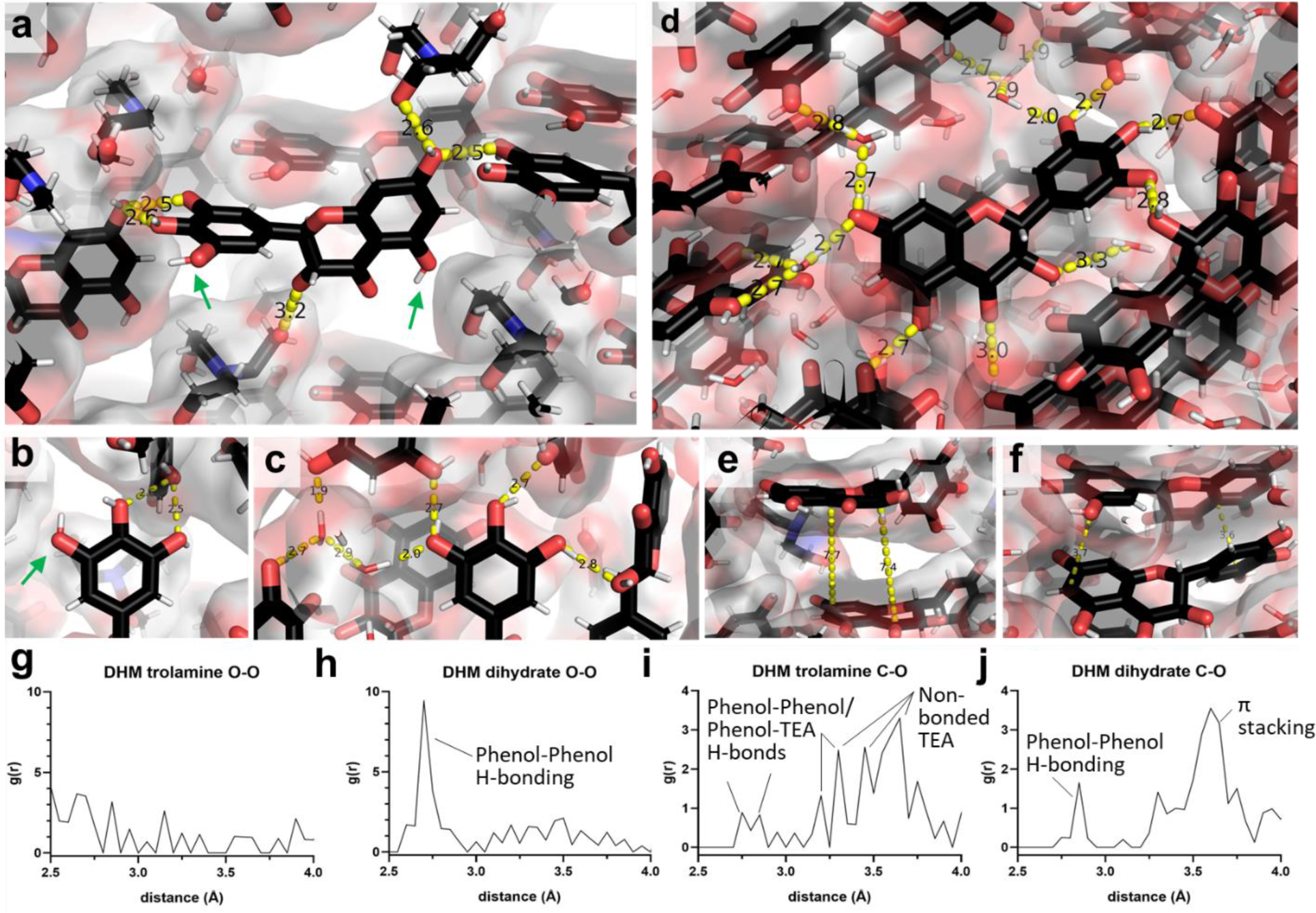
(a-c) DHM-TEA crystal structure. Non-bonded phenols are shown in a and b with green arrows. (d-f) X-ray structure of DHM dihydrate^32^. Contacts between phenols of the pyrogallol ring are shown in b and c for DHM-TEA and DHM dihydrate, respectively. Aromatic stacking interactions are shown in e and f for DHM-TEA and DHM dihydrate, respectively. (g-h) Pair correlation functions for DHM-TEA and DHM dihydrate, as labeled.

Shown in Fig. 5c and 6a, two trolamines pack between DHM molecules disrupting H-bonding between the three phenols of the pyrogallol ring, which are otherwise all H-bonded in the dihydrate crystal (Fig. 6c and d). Trolamine occupies this space without making any H-bond interactions with the pyrogallol, and hence leaves the three phenols of the pyrogallol ring unpaired with gaps of 3.3-3.8 Å between the trolamines and phenols. Moreover, the angles at which the trolamines orient relative to the pyrogallol ring are ∼145-155°, thus leaving space and geometry for water molecules to permeate the lattice of the co-crystal to hydrate and H-bond with the non-paired phenols of the pyrogallol ring.

The number of *in crystallo* phenol H-bonds in DHM-TEA is 5 compared to 9 in DHM dihydrate (Fig. 6d). DHM dihydrate crystals make 5 intermolecular H-bonds between the phenolic moieties of DHM, and an additional 4 water mediated H-bonds bridge neighboring DHM molecules. A single peak at 2.7 Å dominates the O-O pair correlation function, g(r), of the DHM hydrate lattice (Fig. 6h), which reflects the extensive intermolecular network of H-bonded phenols that anneals the O-O pair correlation function. Similarly, a peak at 2.85 Å is in the C-O pair correlation function in Fig. 6j, which arises from intermolecular H-bonding between phenols. Phenols of the DHM-TEA lattice are less extensively H-bonded, with no peak in the O-O pair correlation function (Fig. 6g) and minor peaks in the C-O function 2.75 and 2.85 Å (Fig. 6j). Corresponding, 3 H-bonds are seen between phenols with neighboring DHM molecules and 2 H-bonds with trolamines located on diametric surfaces of DHM (Fig. 6a). The lower overall contact frequency of DHM-TEA in the crystal lattice is consistent with its lower transition temperature measured by DSC for DHM-TEA (Fig. 5e) and explains increased solubility of DHM-TEA. Hydration of DHM-TEA is thermodynamically favored compared with the dihydrate since added enthalpic gains are possible by H-bonds formed between water molecules and non-bonded phenols upon solvation (Fig 6a, green arrows). Solvating DHM dihydrate requires the exchange of intermolecular lattice H-bonds with water molecules, which is kinetically and thermodynamically disfavored since there is no net enthalpy gain to offset the entropic penalty of solvation.

Hydrophobic contacts further contribute to the thermodynamics of DHM solubilization. The C-O pair correlation function shows a peak at 3.7 Å corresponding to the π stacking distances of aromatic rings of DHM in the dihydrate lattice (Fig. 6f and j). The ring-to-ring distance increases to 7.5 Å in the DHM-TEA lattice (Fig. 6e) indicating loss of π stacking. Loss of π stacking is expected to lower the entropic barrier to solvation since aromatic rings of DHM could be partially solvated in the crystal, and moreover the configuration in DHM-TEA eliminates the energetic barrier associated with breaking π stacking in order to solubilize DHM.

## Discussion

Despite tremendous overall H-bonding potential, DHM exhibits remarkable hydrophobic behavior at a macroscopic level. The hydrophobic tendency of DHM is explained by its dihydrate crystal structure, which shows phenols of DHM totally satisfying the H-bonding potentials of neighboring DHM molecules through an extensive network H-bond pairing. In addition, the aromatic rings of DHM form favorable pi-stacking and Van der Waals interactions in the DHM hydrate, resulting in a stable form that is thermodynamically costly to solubilize in water. Our data establishes that trolamine physically disrupts packing of DHM co-crystals and the H-bonding network. These data provide a physical and thermodynamic basis explaining the transformed macroscopic properties of DHM salts, which exhibit up to five times increased water solubility. Co-formers or counterions replace and only satisfy partially the hydrogen bonding potential of DHM, thereby leading to a more favorable thermodynamic state for solubilization. The cryptic hydrogen bonding potential that allows DHM to mask phenols by aggregation may also explain its tendency to better permeate the CNS compared to other polyphenols.

DHM has been investigated for effects in mitigating alcohol use-related disorders, age-related diseases, oxidative stress, poisoning, and liver damage, but its commercial form is a practically water-insoluble solid of beige-brown color. Thus, enhancing the physical properties of DHM to enable improved delivery and biopharmaceutical performance as a pharmaceutically acceptable formulation is of great interest. By behaving as a Brønsted–Lowry acid–base with a log of acid dissociation constant pKa in the useful and relevant pH range of 7-8, we discovered that DHM co-formers or salt formulations are feasible. Ionized species of DHM enable the preparation of a wide range of possible DHM salt forms with bases that satisfy full or partial proton transfer to enable solid and liquid in situ extemporaneous pharmaceutical dosages. As illustrated in Fig. 7, DHM co-crystals described here have appropriate composition to enable improved biopharmaceutical performance in a pharmaceutically acceptable context (e.g. more accurate and efficient dosing of DHM by increasing solubility to allow absorption of higher dose fractions^34^). Regarding the CNS permeability solubility-enhanced DHM, we anticipate circulating DHM could re-establish H-bonding reminiscent the intermolecular phenol-phenol interactions that are seen in the dihydrate crystal form and these assemblies will be important for shielding the hydrophilic character of DHM to enable CNS absorption^35^.

**Figure 7.**
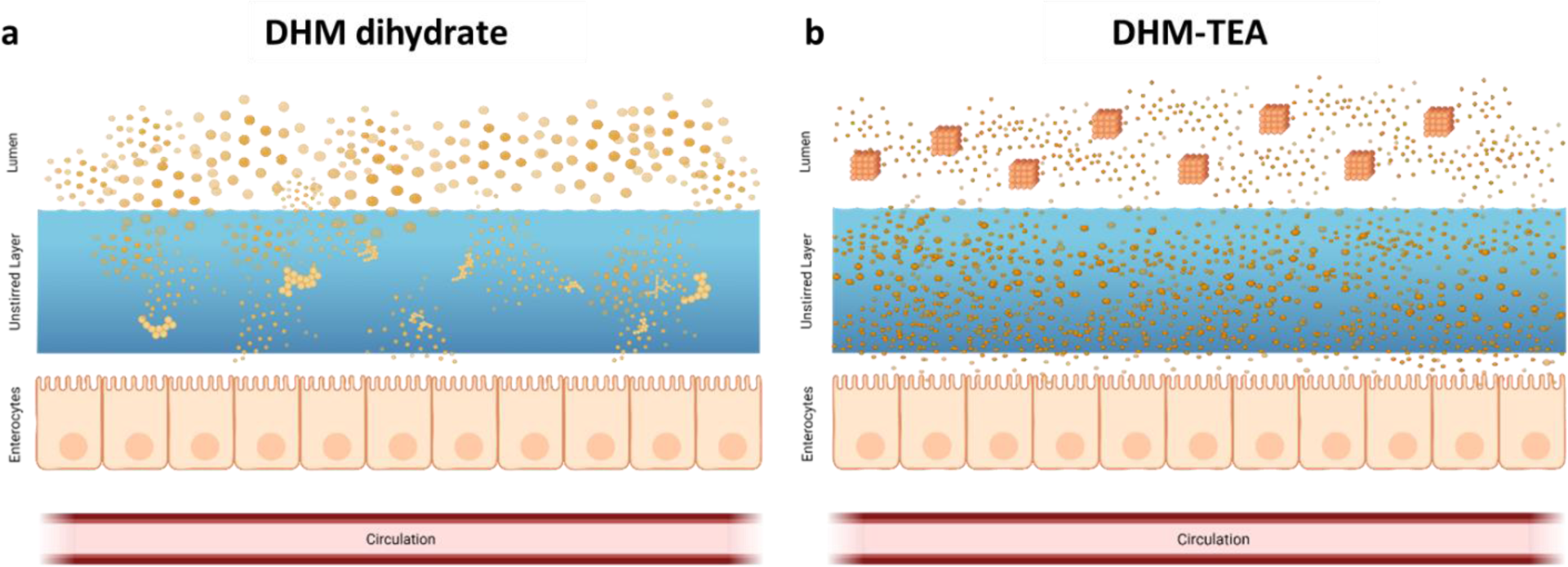
Proposed model for increased DHM delivery by co-former or salt formulation. (a) DHM aggregates resist dissolution and persist in unstirred layer, thereby impeding intestinal absorption. (b) DHM-TEA crystals dissolve more readily in water, thereby increasing fraction of DHM available for absorption, which is beneficial to allow my DHM to reach the systemic circulation.

As a possible tool in the management of Alzheimer’s Disease (AD), DHM holds potential in two possible realms. (1) DHM is a lead for medicinal chemistry structure-activity studies to generate pharmaceutical-grade anti-tau medications with improved potency and CNS permeability. (2) DHM natural products as dietary supplements offer a more immediate possible approach to combine with other lifestyle adaptations to aid healthier aging. Natural products are affordable and readily accessible chemicals, and in the case of DHM, could be rapidly developed and deployed commercially to support studies investigating healthier aging alongside lifestyle modifications (i.e. attention to diet, exercise, and cardiovascular health). The MicroED structure shows that co-crystalline formulations of DHM remain unaltered in chemical structure despite improvements in aqueous solubility and apparent potency. Thus, the formulations we describe do not alter the natural product status of DHM. DHM is sold as a dietary supplement in the United States and is generally regarded as safe (GRAS) by the FDA. A human clinical trial of 60 adults with non-alcoholic fatty liver disease assigned DHM 150 mg twice daily vs. placebo for three months identified no significant safety concerns.^36^ Although, as with most flavonoids, the major obstacle to DHM’s clinical application is its suboptimal pharmacokinetics (PK). We anticipate the new formulations of DHM described here with greater solubility will lend to improved bioavailability, although confirmation remains to be seen.

In summary, by strategically coupling structural and formulations chemistry, DHM was realized as a possible natural product inhibitor lead compound for targeting prionogenic seeding by AD tau. These studies open the door for near-term investigations of the effects of DHM salts as dietary supplements for testing in humans to support healthier ageing, and in the broader context, suggests co-crystalline and/or salt forms may be extended to formulate other pseudo-hydrophobic medicinal chemicals to enhance solubility and efficiencies of dosing.

## Materials & Methods

A commercially available batch of dihydromyricetin [(DHM) IUPAC Name (2R,3R)-3,5,7-trihydroxy-2-(3,4,5-trihydroxyphenyl)-2,3-dihydrochromen-4-one, of ≥ 98% purity] was purchased from Master Herbs Inc (Pomona, CA). Counterions, triethanolamine (99%) was purchased from Lab Alley (Spicewood, TX); sodium hydroxide, calcium hydroxide, and L-Lysine were all purchased from Sigma-Aldrich (Burlington, MA); and TRIS-base was purchased from Gold Biotechnology (St. Louis, MO). Solvents, DMSO, pure ethyl alcohol (200 proof, molecular grade), methanol (≥99.5% purity, molecular grade), 2-propanol (≥99.5% purity, molecular grade), acetone (≥99.5% purity, molecular grade), and anti-solvent, chloroform (anhydrous, ≥99%, containing 0.5-1.0% ethanol as stabilizer) were all purchased from Sigma-Aldrich (Burlington, MA).

The instruments used were a Mettler-Toledo ME54TE/00 analytical balance and TE412 Top-loading balance (Switzerland), VWR sypHony B10P pH meter, Fischer Scientific Stirring Hotplate 11-500-49SH (Walthamm MA); magnetic stir bars, flat bottom 96-well plates, 20mL glass scintillation vials, Vacubrand ME 2 NT vacuum pump, and the Biotek Agilent Synergy HTX Multi-mode Reader (Santa Clara, CA); qualitative Whatman Filter Papers (47mm diameter) was purchased from Sigma-Aldrich (Burlington, MA). Polarized light microscopy was performed using Olympus BX51 Microscope, and the analyzer (U-ANT; Analyzer for transmitted light U-P115) and polarizing (U-POT; Polarizer for transmitted light, 45mm U-P110) filters were purchased from Olympus (Waltham, MA). Melting point determinations were performed using PerkinElmer DSC 8500 with HyperDSC and cooling accessory (IntraCooler 2), (PerkinElmer, Inc., Valencia, CA). X-ray powder diffraction studies were performed using MiniFlex™ 600, (Rigaku, Inc., The Woodlands, TX).

### Counterion Screen

Various counterions indicated in the main text were screened by preparation in a variety of solvents (methanol, ethanol, 2-propanol, or acetone) to facilitate crystallization with DHM by slow evaporation. Crystallization was carried out at ambient temperature (22-25°C) in 250mL Erlenmeyer flasks. A 1:1, 1:1.25, or 1:2 molar ratio of DHM to counterion were tested to find optimum stoichiometry for crystallization. The counterion was first dissolved in the respective solvent and DHM added and to it with stirring for at least 45 min until homogenous slurries were observed. Slurries were then shielded from light and allowed to slowly evaporate for two weeks. Resulting powders were washed with 50mL of chloroform over a vacuum pump membrane filtration setup. Filtrate residue was dried, recovered from filter membrane, and stored in sealed glass vials at ambient conditions further characterization and biological activity testing.

### Polarized Light Microscopy

An Olympus (Olympus, Waltham, MA) microscope equipped with a polarizer and analyzer was used to initially evaluate isolated solids from slurries. 1000x magnification was used with an analyzer (U-ANT) slider insert and a polarizer (U-POT). Samples were qualitatively assessed for particle size and shape, as well as birefringence for initial evidence of existence of orientation-dependent differences in refractive index.

### pKa Determination

pKa values were calculated using ADMET Predictor™ version 10.3.0.7 64-bit edition module of Simulations Plus software (Lancaster, CA), and ChemDraw® Professional version 20.1.1.125 (PerkinElmer Informatics, Inc.). Potentiometric titration curves of DHM were obtained by adding 0.1mL of a 0.5N NaOH standard solution to three different 2mg/mL DHM analyte solutions. All solutions were made in 50mL beakers using deionized water and prepared at 22-25°C. The 2mg/mL DHM standard solution was titrated with 0.5N NaOH. The NaOH standard solution was prepared by dissolving 3g of sodium hydroxide in 150mL of water for a final concentration of 0.5N NaOH. A 100mg/mL stock solution of DHM was prepared by weighing 0.5g of DHM using an analytical balance and dissolving it in 5mL of methanol in a 15mL conical tube. Three analyte solutions were prepared using methanol and water mixtures to yield 25mL of solutions with 25, 50, and 75% v/v methanol content. Five hundred µL of DHM stock solution was then added to the three analyte solutions to yield final DHM concentration of 2mg/mL, the initial pH value was measured and recorded.

pKa was orthogonally determined through UV-VIS spectroscopy optical absorption technique described by Pandey, et. al.^37^ that was adapted and modified. Briefly, a 5mM analyte stock solution of DHM was prepared by measuring 16 mg of DHM dissolved in 10mL of DMSO and was further diluted to 2mM and used as the working stock solution. Buffer solutions of various pH values were prepared as described in Table 1. Using a 96-well plate, 0.2mL of each pH buffer solution was added to duplicate wells, then 0.5μL/well of the DHM stock solution was added for a final DHM concentration of 5μM in each sample well. A spectral scan was conducted (λ = 230-700nm), and absorbance was measured in 10nm increments. The pKa was calculated using pKa=pH+log [(DHM^0^ -DHM^pH^)/(DHM^pH^ -DHM^-^), where DHM^0^ is the absorbance of the unionized molecule at lowest pH, DHM^pH^_abs_ is the absorbance of the molecule in respective buffers tested, and DHM^-^ is the absorbance of the ionized molecule at highest pH.

### Differential Scanning Calorimetry

Samples (3-5mg, small enough to avoid saturating the apparatus) were analyzed using a PerkinElmer DSC 8500, (PerkinElmer, Inc., Waltham, MA), that is equipped with a cooling system that uses a continuous dry nitrogen purge at 25mL/min. The instrument was calibrated for temperature and enthalpy change with indium, tin, and lead samples provided by the manufacturer. The samples and reference (i.e., blank) were placed in the furnace. Data was collected from 25-300 °C with a heating rate of 10 °C/min. Onset and peak of observed thermal events were recorded and qualitatively analyzed for relative comparison.

### X-Ray Powder Diffractometry

Solid material isolated from DHM slurries were broken down into a fine powder using a glass coverslip or a spatula. Approximately 5-10 mg of the powder was placed onto a zero background, 5mm diameter × 0.2mm deep well, silicone sample holder (Rigaku, Inc., The Woodlands, TX). Analysis was performed on a Rigaku Miniflex 600 Benchtop XRD System (Rigaku, Inc., The Woodlands, TX). Diffractograms were collected operating at a scanning rate of 2.00° min^-1^. The diffraction spectra were recorded at the diffraction angle, 2θ from 0° to 50° at room temperature.

### In silico modeling

Although DHM has a similar chemical structure to EGCG (Fig. 1a), the binding site of DHM remains unknown. Therefore, molecular docking of DHM on paired helical filament (PHF) tau (PDB: 5O3L) was performed to uncover potential sites of binding using Cavity-detection guided Blind Docking (CB-Dock), a protein-ligand docking method that uses AutoDock Vina, developed by the Yang Cao Lab at Sichuan University (Fig. 1b). (Yang Cao and Lei Li. Improved protein-ligand binding affinity prediction by using a curvature-dependent surface-area model,. Bioinformatics, 2014). The top five potential binding cavities were identified.

### Preparation of crude Alzheimer’s brain-derived tau seeds

Human Alzheimer’s brain autopsy samples were obtained from the UCLA Pathology Department according to HHS regulation from patients who consented to autopsy. Approximately 0.2 g of hippocampal tissue was excised, and a Kinematica PT 10-35 POLYTRON was used to homogenize the tissue with sucrose buffer supplemented with 1 mM ethylene glycol tetraacetic acid (EGTA) and 5 mM Ethylenediaminetetraacetic acid (EDTA) at level 4-5 in 15 ml falcon tubes.

### K18CY cell culture (find reference to summarize)

HEK293T cell lines that stably express tau-K18CY labeled with green fluorescent protein (GFP) obtained from Marc Diamond’s laboratory at the University of Texas Southwestern Medical Center (Sanders et al., 2018) were used. The cells were cultured in a T25 flask in Dulbecco’s Modified Eagle Medium (DMEM) (Life Technologies, cat. 11965092) supplemented with 10% (vol/vol) Fetal Bovine Serum (FBS) (Life Technologies, cat. A3160401), 1% penicillin/streptomycin (Life Technologies, cat. 15140122), and 1% Glutamax (Life Technologies, cat. 35050061) at 37°C and 5% CO_2_ in a humidified incubator. To test the inhibitors on the biosensor cells, 100 µl of cells were plated 1:10 in 96 well plates and stored in the 37°C, 5% CO_2_ incubator for 16 to 24 hours prior to transfection.

### Screening of DHM alongside EGCG

EGCG and DHM were dissolved in dimethyl sulfoxide (DMSO) to 10 mM at room temperature. Previously homogenized human-derived AD crude brain extracts were diluted 1 to 20 with Opti-MEM (Thermo Fisher Scientific, cat. 31985062) and sonicated in a Qsonica multiplate horn water bath for 3 minutes at 40% power. The diluted brain extracts were then incubated with the inhibitors for 16 to 24 hours at 4°C to yield a final EGCG or DHM concentration of 10 mM on the tau K18CY biosensor cells. Inhibitor-treated seeds were sonicated again in a Cup Horn (manufacturer?) water bath for 3 minutes at 40% power and then mixed with a 1 to 20 solution of Lipofectamine 2000 (Thermo Fisher Scientific, cat. 11668019) and Opti-MEM. The Lipofectamine creates a liposome around the fibrils to allow delivery into the cells. After 20 minutes, 10 µl of inhibitor-treated fibrils were added to the previously plated 100 µl of cells in triplicate, avoiding use of the perimeter wells.

### Screening of DHM crystalline co-formers or salts

DHM and five co-former or salt formulations were screened alongside EGCG in triplicate using the same tau prionogenic seeding assay workflow, except inhibitors were dissolved in sterile, deionized water rather than DMSO, at 37°C or on the benchtop at room temperature. Dissolved inhibitors were then centrifuged for 10 minutes at 8,000 or 15,000 rpm to pellet any insoluble fraction. The supernatants were removed and used to test the inhibitory effect of the soluble fraction of aqueous DHM which resulted from co-former or salt form dissolution.

### Plate reading

The number of seeded aggregates was determined by using the BioTek Cytation 5 Imaging Multimode Reader in the GFP channel to image the entire 96 well plate. A 3×2 montage was used to capture as much area of each well as possible. Exposure and contrast were adjusted to allow puncta to distinguish the seeded aggregates from the cells. Seeded aggregates appear as bright green puncta (Fig. 1c)

### Quantification of tau aggregation

Seeded aggregates in a given image were quantified using a Java-based image processing program ImageJ 2.3.0 (Schneider et al., 2012)) script, which subtracts the background fluorescence from unseeded cells, and then uses a built-in particle analyzer to quantify the number of puncta (seeded aggregates) as peaks with fluorescence that contrast the background. The puncta count was then normalized across all images according to cell confluence by using a separate ImageJ 2.3.0 script for confluence. Normalizing to confluence is necessary in the event that an inhibitor is toxic to cells, so that toxic inhibitors are not considered to be effective against tau seeding. The average number of seeded aggregates of each well and standard deviations from triplicate measurements, normalized to confluence, was plotted to compare inhibitory activity.

### Determination of potencies

The same tau prionogenic seeding assay workflow with Alzheimer’s crude brain extracts was conducted to determine the IC_50_’s – half maximal inhibition – of EGCG, DHM, and DHM analog. Ten different final inhibitor concentrations were tested – 0, 0.1, 0.25, 0.5, 1.0, 2.5, 5.0, 10.0, 20.0, and 40.0 µM – in triplicate. Inhibitors were dissolved in both DMSO and DI water. After aggregation in each well was quantified using ImageJ, the GraphPad Prism 9 software was used to generate dose-response curves for IC_50_ determination.

### Determination of solubility

Solubility determinations of DHM and salt or co-crystal formulations were conducted by spectrophotometric analysis. Inhibitors were first dissolved to 10 mM in DMSO, and the absorbance spectra were collected to determine the wavelength at which absorbance peaked. The 10 mM stocks were then diluted 1:3, 1:6, 1:12, 1:30, 1:60, 1:120, and 1:300 and absorbance of each was measured using the NanoDrop One/OneC Microvolume UV-Vis Spectrophotometer at peak wavelength alongside a DI water negative control to produce a calibration curve. The working linear range was determined by eliminating absorbance measurements above the limit of linearity. The 10 mM stocks of DHM and salt co-crystal forms previously used for the tau prionogenic seeding assays were diluted as necessary to fall within the linear range and their measured absorbances were used to calculate the true concentrations of stocks solutions.

### Transmission electron microscopy (TEM)

Transmission electron microscopy was used to determine whether DHM crystalline co-former or salt formulations exert inhibitor effects on seeding through tau disaggregation like EGCG. Purified Alzheimer’s brain-derived tau fibrils were incubated with inhibitor and added to negative stain grids to be imaged with the JEOL-2100 TEM.

### Preparation of purified Alzheimer’s brain-derived tau fibrils

The same homogenization process of human Alzheimer’s brain autopsy samples with sucrose buffer supplemented with 1 mM EGTA and 5 mM EDTA to prepare the crude brain extracts was conducted. Protein content was then precipitated by heating in the Biorad thermal cycler to 95°C for 20 minutes. Precipitated protein homogenates were then combined and centrifuged at 20,100 × g for 30 minutes at 4°C in an Eppendorf centrifuge, and the resulting supernatants were transferred to airfuge tubes and ultracentrifuged at 95K for one hour. Ultrapellets containing the purified fibrils were resuspended in 1X phosphate buffered saline (PBS), pH 7.4.

### Negative stain grid preparation

Purified Alzheimer’s brain-derived tau fibrils were diluted 1:10 in PBS and incubated with EGCG or DHM ligands for 48 hours at 4°C. Negatively stained EM grids were prepared by depositing 6μl of fibril samples on formvar/carbon-coated copper grids (400 mesh) for 3 minutes with inhibitor pre-incubation times of either 0 hours (negative control) or 48 hours (positive control). The sample was rapidly and carefully removed by fast blot using filter paper without drying the grid and stained with 4% uranyl acetate for 2 minutes, then wicked dry by filter paper.

### Quantitative EM (qEM) imaging

For quantitative EM image (qEM), negatively stained EM grids of each sample were screened on the JEOL 2100 TEM at a magnification of ×12,000, collecting 99 images in consistent increments. Visible fibrils were counted manually and analyzed in triplicate groups of 33 micrographs for each experimental condition.

### MicroED sample preparation, data collection and processing

The DHM-TEA co-crystal was prepared for MicroED as described previously.^38^ Around 1 mg ground powder was transferred into a 10mL scintillation vial and mixed with a carbon-coated copper grids (400-mesh, 3.05 mm O.D., Ted Pella Inc.) which was pretreated with glow-discharge plasma at 15 mA for 60 s on the negative mode using PELCO easiGlow (Ted Pella Inc.). After a gentle shaking of the vial, the grid was taken out and clipped at room temperature.

The clipped grid was loaded in an aligned Thermo Fisher Talos Arctica Cryo-TEM (200 kV, ∼0.0251 Å) at 100 K, equipped with a CetaD CMOS camera (4096 × 4096 pixels). Screening of size-suitable microcrystals was done in the imaging mode (SA 3400×). The MicroED data was collected in the diffraction mode with 829 mm diffraction length, 70µm C2 aperture and a 100µm selected area aperture in the parallel beam condition (45.2% C2 intensity) which resulted a beam size at approximately 2.5 µm. Typical data collection was performed using a constant rotation rate of ∼1 deg/s over an angular wedge of 130° from -65° to +65°, with 1s exposure time per frame. Crystals as selected for MicroED data collection were isolated and calibrated to eucentric height to maintain the crystal inside the beam during the rotation.

The MicroED data was saved in MRC format and converted to SMV format using the mrc2smv software (https://cryoem.ucla.edu/microed). The converted frames were indexed and integrated by XDS^39^. Then two datasets were scaled and merged using XSCALE^39, 40^, and the intensities were converted to SHELX hkl format using XDSCONV^39, 40^. The merged dataset can be *ab initio* solved by SHELXT^41^ and refined by SHELXL^42^ to yield the final MicroED structure.

## Acknowledgements

We thank the staff at the USC Center for Nanoimaging and the UCLA MicroED Center for supporting EM work (National Institutes of Health P41GM136508). This work was supported in part by the Timothy M. Chan Professorship in Complementary Therapeutics (USC) to DLD, NIH 1R01AG070895-01A1 (DSE), NIAAA R01AA022448 (DLD), USC Institute of Addiction Science (IJU), and the University of Southern California Summer Undergraduate Research Fund (SURF). The Gonen laboratory is supported by funds from the Howard Hughes Medical Institute. Select images were created by Biorender.com. Data was deposited to CCDC accession number XXXX.

## Supplementary Figures

**Supplementary Figure 1.**
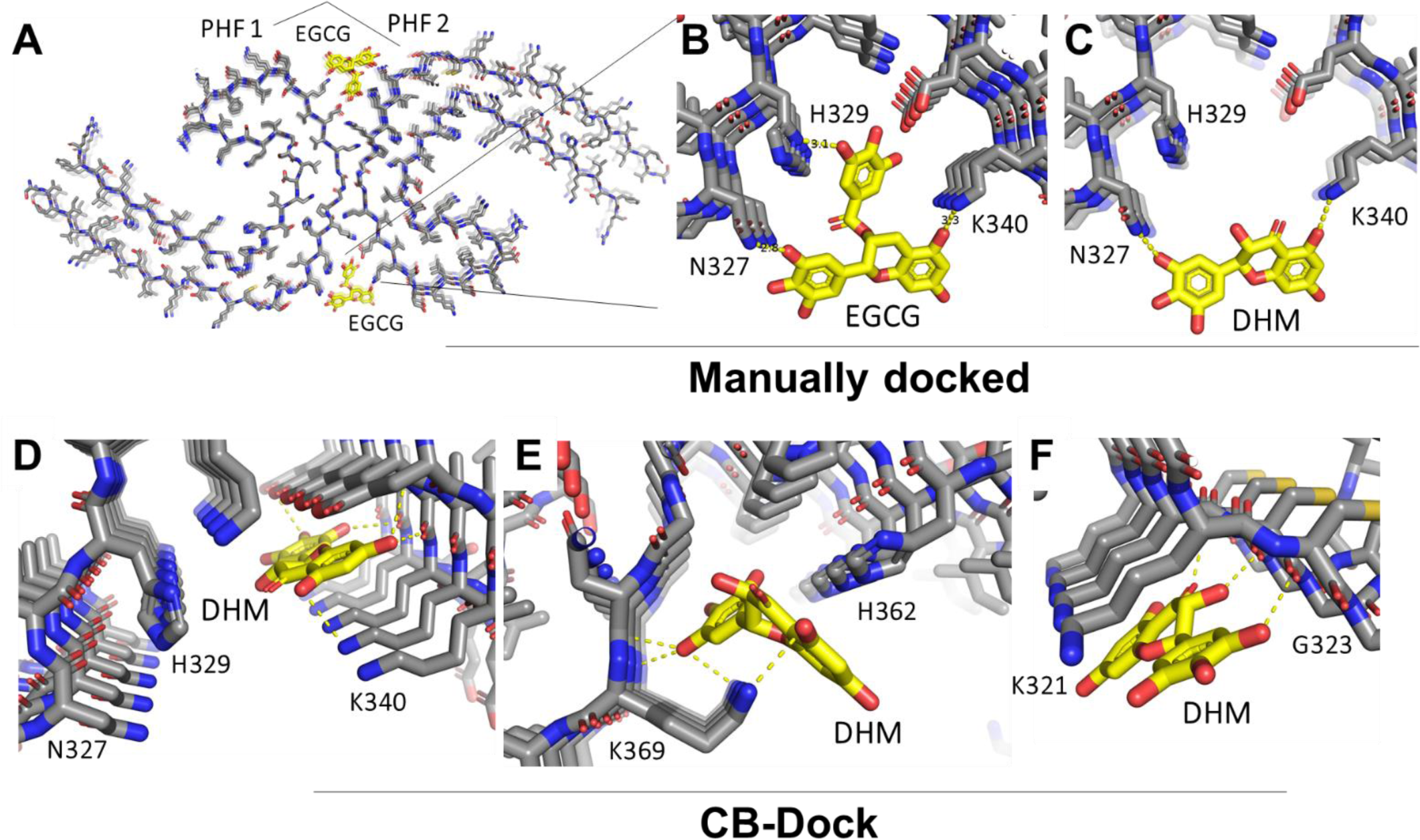
(A and B) CryoEM structure of AD brain-derived tau PHF bound by EGCG. (C) Manually generated model of DHM binding to EGCG binding site of AD tau made by superimposing DHM with bound EGCG. (D-F) Alternative predicted DHM binding sites generated by CB-Dock. (D) Site 1 overlaps with the binding volume predicted in the manually generated model in C, except bound DHM is predicted to orient with aromatic rings perpendicular to the fibril axis burying in a cavity between Lys340 and Glu338. (E) Site 2 of DHM binding predicted by CB-Dock occurs in a cavity lined by His362 and K369. (F) Site 3 predicted by CB-Dock occurs with DHM packing aromatic moieties with the aliphatic chain of K321, and H-bonding occurring between the phenolic moieties and the amide backbone at a sharp turn formed by G323.

**Supplementary Figure 2.**
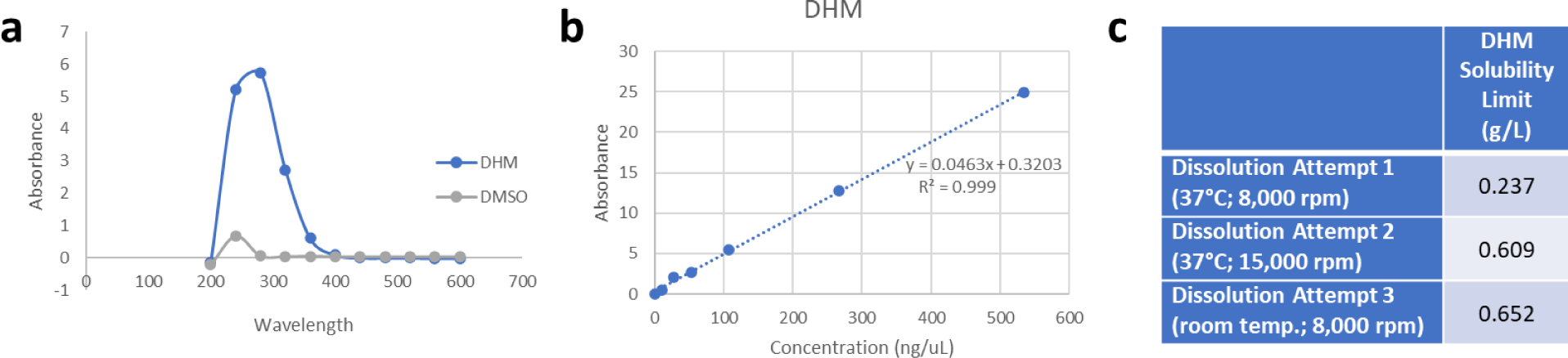
Aqueous solubility determination of DHM. (a) Absorbance scan of DHM dissolved in DMSO shows a peak at 280 nm. (b) Standard curves of DHM supernatants dissolved in water to a target concentration 10 mM and clarified by centrifugation. (c) Table detailing concentrations measured from DHM supernatants dissolved in water and determined from standard curve in b.

**Supplementary Figure 3.**
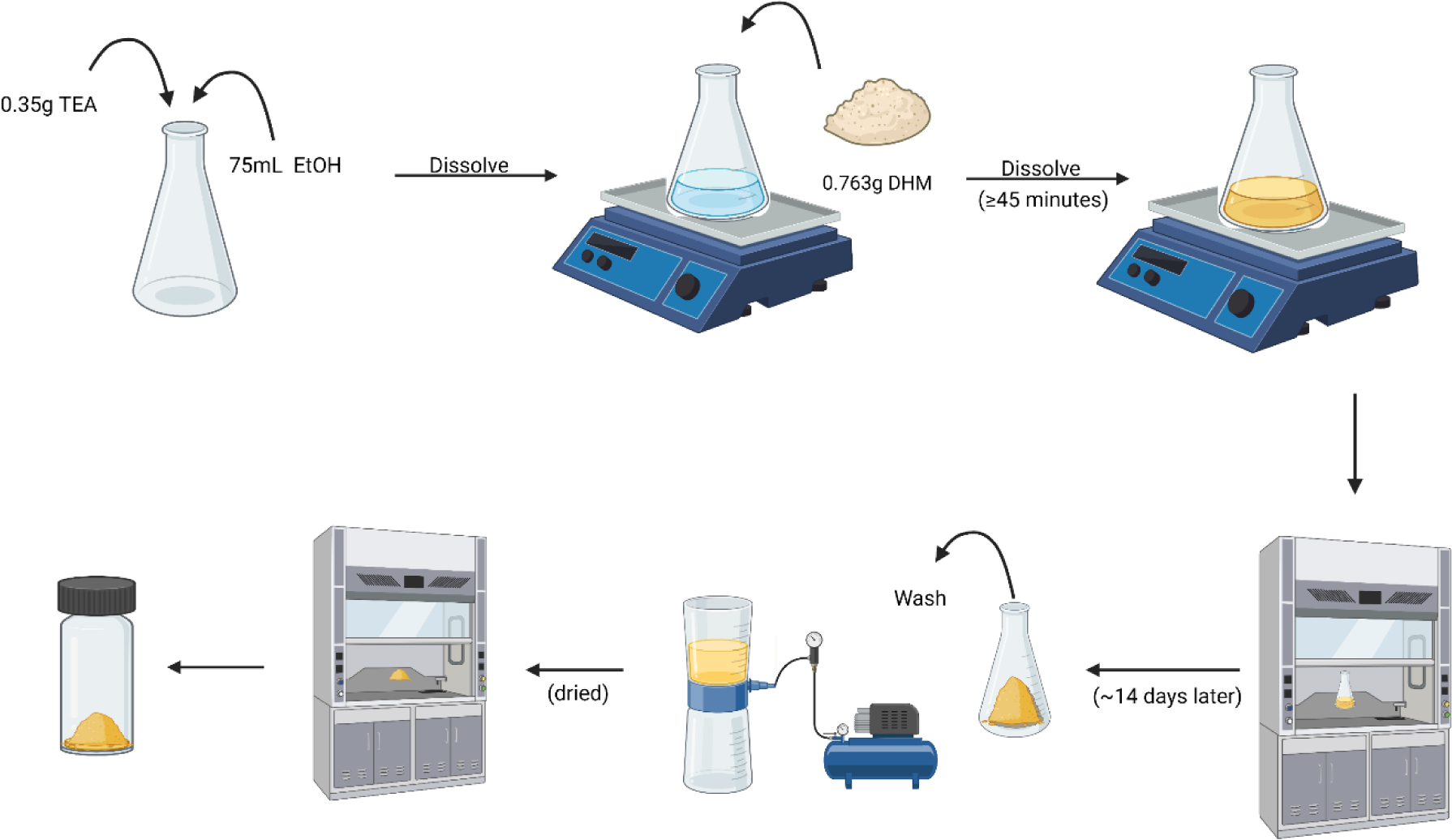
Example of a slurry preparation using DHM and TEA. As shown, mixture was achieved by preparing a solution containing 0.35g of TEA dissolved in 75mL of ethyl alcohol (200 proof) on a magnetic stir plate. Once dissolved, 0.763g of DHM powder (1:1 molar ratio) was added to the solution containing TEA and stirred for at least 45 minutes. Precipitate recovered 14 days later was placed in a membrane filtration apparatus and vacuum washed with chloroform to remove possible remaining excess chloroform-extractable material, and then dried. After drying, solids were stored in a clear glass scintillation vial at room temperature (22-27°C) until future use.

**Supplementary Figure 4.**
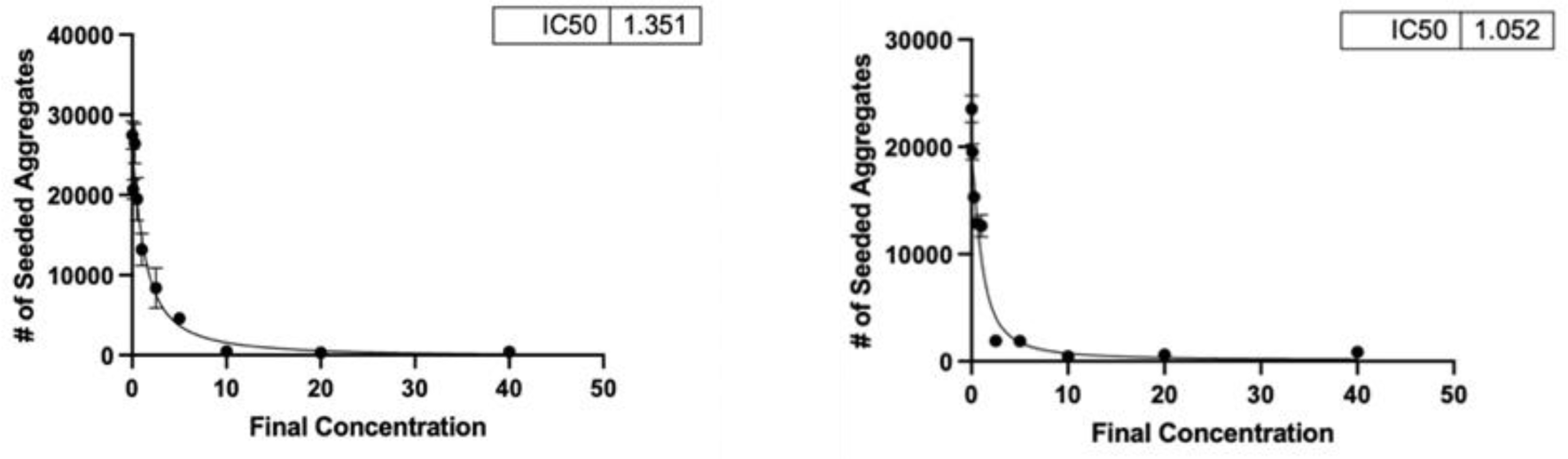
IC50 curves generated as described in the main text, except using DHM-TEA and -Ca(b) formulations (left and right, respectively) dissolved in DMSO. Both formulations have IC50s similar to DHM when dissolved in DMSO (Fig. 1B main text), thus suggesting DHM is the active component in tau inhibitor assays and that counterions themselves do not exhibit inhibitory activity towards tau seeding.

**Supplementary Figure 5.**
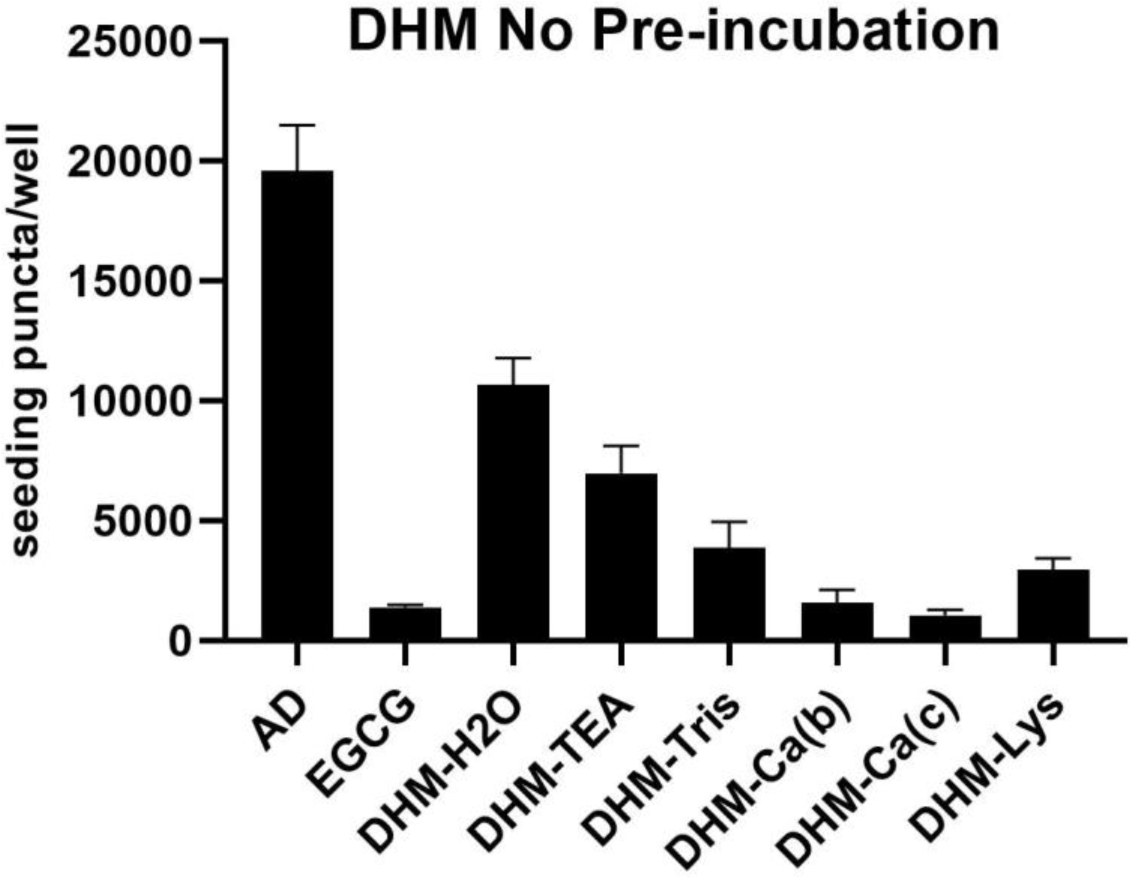
Seeding inhibition measured as described in the main text, without inhibitor pre-incubation with crude AD brain homogenates.

**Supplementary Figure 6.**
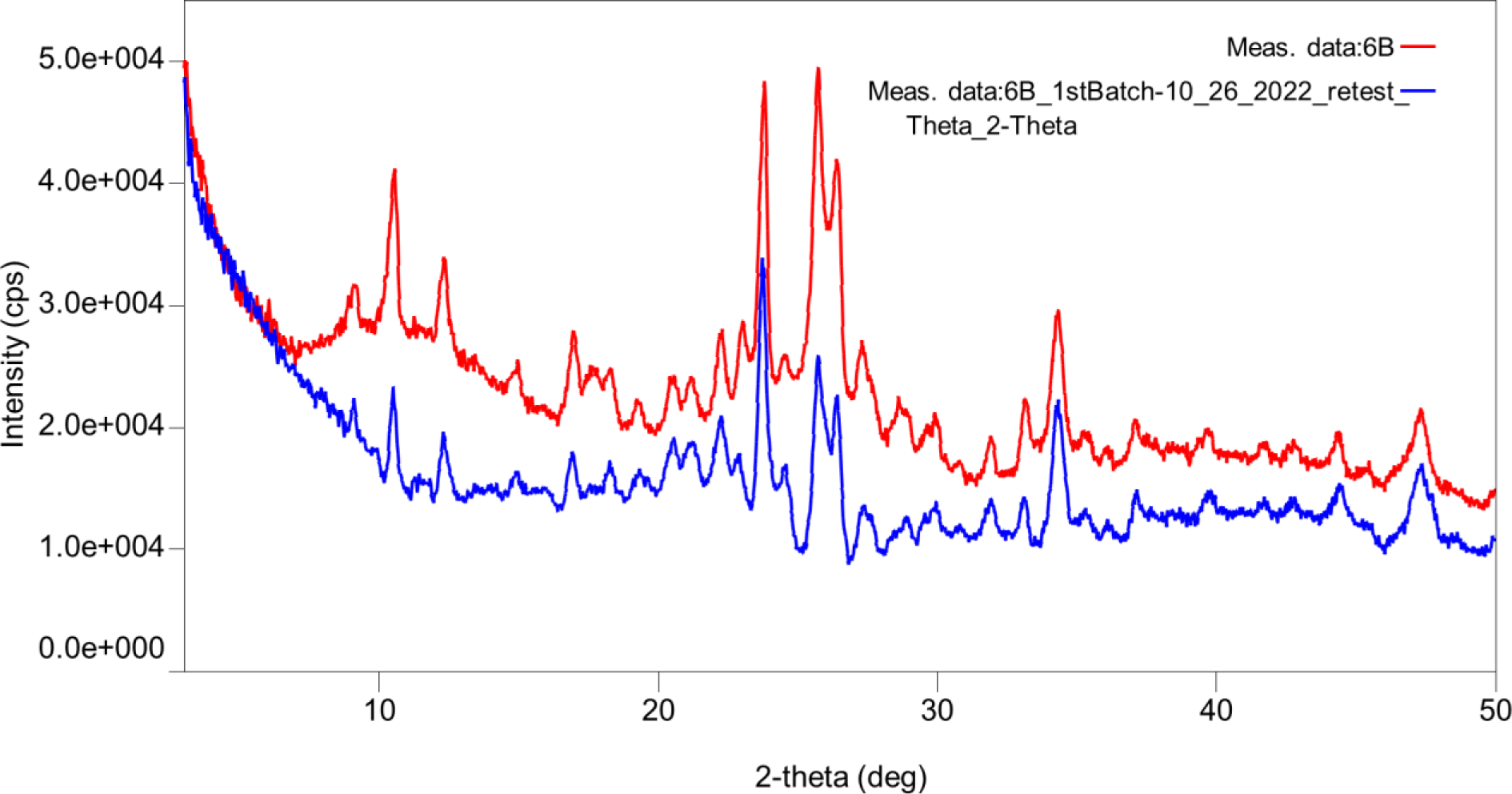
X-Ray powder diffractogram comparing DHM-Ca salt b freshly prepared (red) overlayed with the same preparation aged 1 year at room temperature (blue).

**Supplementary Figure 7.**
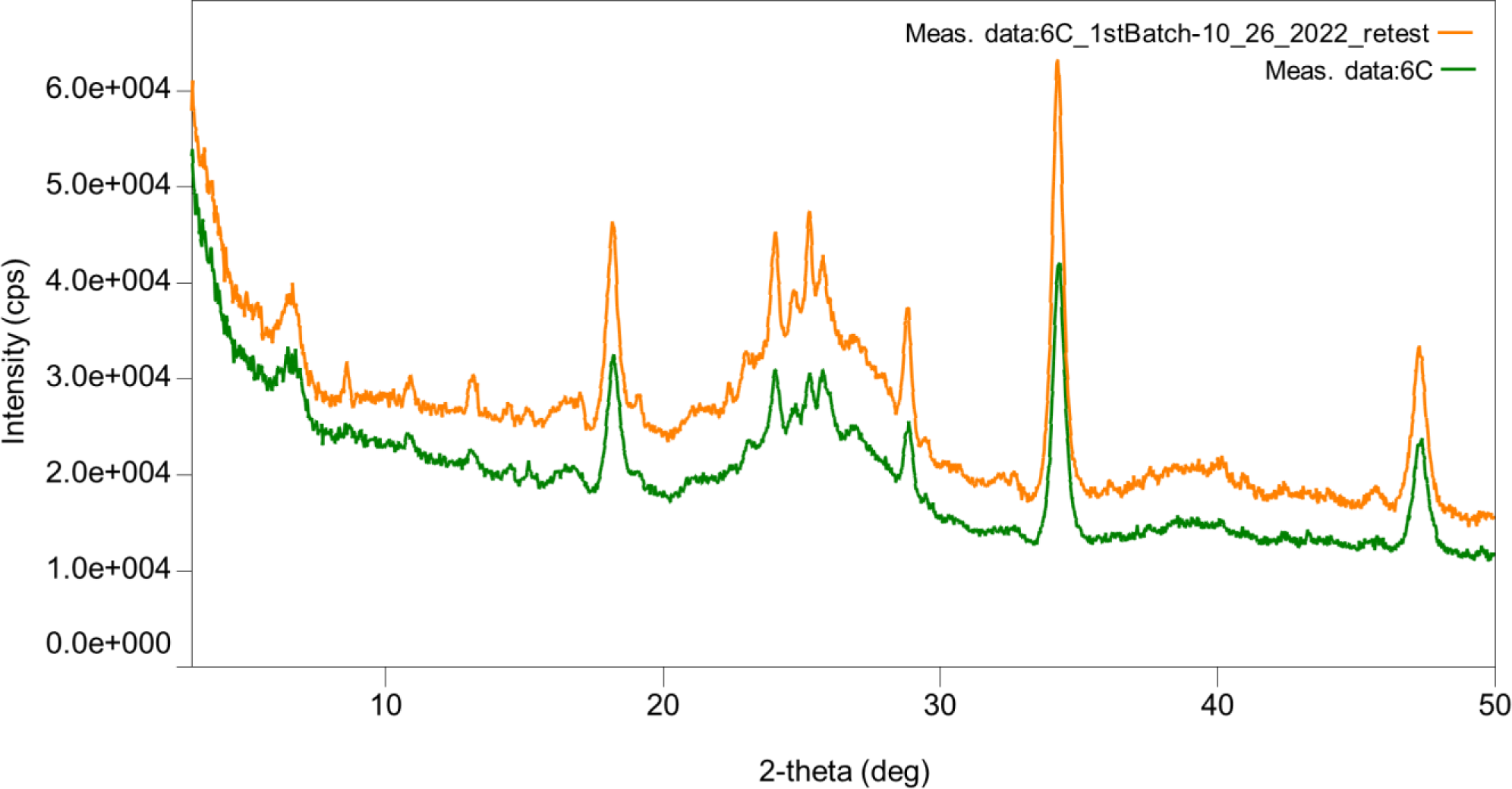
X-Ray powder diffractogram comparing DHM-Ca salt c freshly prepared (orange) overlayed with the same preparation aged 1 year at room temperature (green).

**Supplementary Figure 8.**
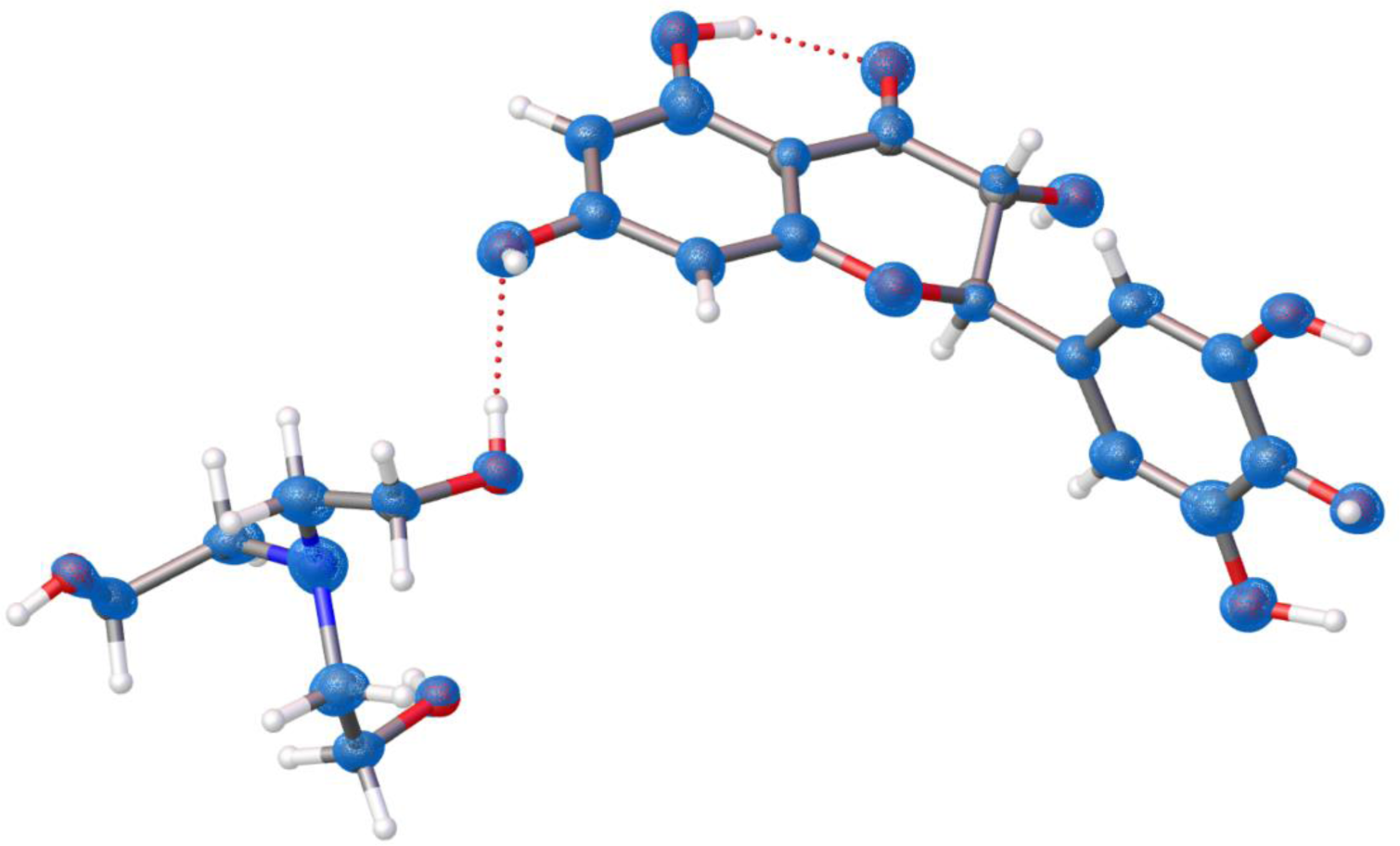
The crystal structure of DHM-TEA as solved with a resolution of 0.75 Å. The blue mesh are 2Fo-Fc electrostatic potential maps at the level of 0.67 e·Å-3.

## Supplementary Tables

**Table 1.** List of pH buffers.

| Buffer Preparation (in 80mL H<sub>2</sub>O) |  |  |  |  |  |  |  |  |
| --- | --- | --- | --- | --- | --- | --- | --- | --- |
| pH | 3.1 | 4.3 | 5.8 | 7.5 | 8.4 | 9.4 | 10.3 | 13.3 |
| Components and q.s. | 0.48g Citric Acid<br>0.81g NaOH<br>0.51g HCl<br>Adjust to final pH, add water to 100mL | 0.48g Citric Acid<br>0.81g NaOH<br>0.2g HCl<br>Adjust to final pH, add water to 100mL | 0.48g Citric Acid<br>1.8g NaOH<br>0.46g HCl<br>Adjust to final pH, add water to 100mL | 0.3g Tris-HCl<br>0.07g HCl<br>Adjust to final pH, add water to 100mL | 0.3g Tris-HCl<br>0.03g HCl<br>Adjust to final pH, add water to 100mL | 0.214g Na<sub>2</sub>CO<sub>3</sub><br>1.17g NaHCO<sub>3</sub><br>Adjust to final pH, add water to 100mL | 187.4mg Na<sub>2</sub>CO<sub>3</sub><br>201.6mg NaHCO<sub>3</sub><br>Adjust to final pH, add water to 100mL | 1.6g NaOH (0.05N NaOH) |

**Table 2.** List of counterions and solvents used for counterion screen study.

| Base/Counterion | Solvent |
| --- | --- |
| Triethanolamine (TEA)<br>pKa: 7.74<br>(MW: 149.188 g/mol) | Ethanol (A) |
|  | Methanol (B) |
|  | 2-propanol (C) |
|  | Acetone (D) |
| Sodium Hydroxide (NaOH)<br>pKa: 15.7<br>(MW: 39.99 g/mol) | Ethanol (A) |
|  | Methanol (B) |
|  | 2-propanol (C) |
|  | Acetone (D) |
| TRIS Base (2-Amino-2-(hydroxymethyl)-1,3-propanediol)<br>pKa: 8.1<br>(MW: 121.14 g/mol) | Ethanol (A) |
|  | Methanol (B) |
|  | 2-propanol (C) |
|  | Acetone (D) |
| L-Lysine<br>pKa: 10.53<br>(MW: 146.19 g/mol) | Ethanol (A) |
|  | Methanol (B) |
|  | 2-propanol (C) |
|  | Acetone (D) |
| Calcium Hydroxide (Ca (OH)<sub>2</sub>)<br>pKa: 15.7<br>(MW: 74.09 g/mol) | Ethanol (A) |
|  | Methanol (B) |
|  | 2-propanol (C) |
|  | Acetone (D) |

**Table 3.** MicroED data processing statistics of DHM-TEA.

|  |  |
| --- | --- |
| Stoichiometric formula | C<sub>21</sub> H<sub>27</sub> N O<sub>11</sub> |
| Mr | 469.44 |
| Temperature (K) | 100 |
| Crystal system, space group | Triclinic, P-1 |
| Unit cell lengths a, b, c (Å) | a 8.0(10) b 11.1(10) c 12.2(10) |
| angles $\alpha$, $\beta$, $\gamma$ (°) | $\alpha$ 98.1(10) $\beta$ 101.4(10) $\gamma$ 106.4(10) |
| Cell volume (Å<sup>3</sup>) | 996.241 |
| Overserved reflections (#) | 10956 |
| Unique reflections (#) | 4415 |
| R<sub>obs</sub> (%) | 14.5 |
| R<sub>meas</sub> (%) | 18.5 |
| CC1/2 | 98.9 |
| Resolution (Å) | 0.74 |
| Completeness (%) | 86.7 |
| R<sub>1</sub> (%) | 19.5 |
| wR<sub>2</sub> (%) | 46.6 |
| GooF | 1.562 |

